# PRECOG update: An augmented resource of clinical outcome associations with gene expression for pediatric and immunotherapy cohorts

**DOI:** 10.1101/2025.08.22.671849

**Authors:** Brooks A. Benard, Chinmay K. Lalgudi, Ilayda Ilerten, Ruohan Wang, Andrew J. Gentles

## Abstract

Gene expression can be used to define prognostic and predictive biomarkers across cancers and treatment modalities. PRECOG (https://precog.stanford.edu) is a compendium of datasets with gene expression and clinical outcomes that facilitates visualization of associations between genomic profiles and patient survival. Here, we augment the existing PRECOG with new datasets in previously poorly represented adult cancer types, as well as adding annotated pediatric and immunotherapy treated cohorts. Pediatric PRECOG comprises ∼4,000 patients across 12 cancers; while the immunotherapy cohort (ICI PRECOG) contains ∼4,500 patients across 20 cancer subtypes from 80 distinct datasets across 52 studies. We compute and visualize associations of gene expression with survival outcomes using Cox regression for time-to-event, or logistic regression for responder vs non-responder, across all datasets. We also estimate cell type fractions in samples via computational deconvolution using CIBERSORTx, to identify survival associations at the level of cell types. All expression data, clinical annotations, and gene and cell type survival z-scores and meta z-scores for adult, pediatric, and ICI PRECOG, are available for interactive analysis and download, along with Kaplan-Meier and boxplot visualizations. This updated resource will provide new insights into biomarkers for specific therapies, populations, and cancer types.

**GRAPHICAL ABSTRACT:** 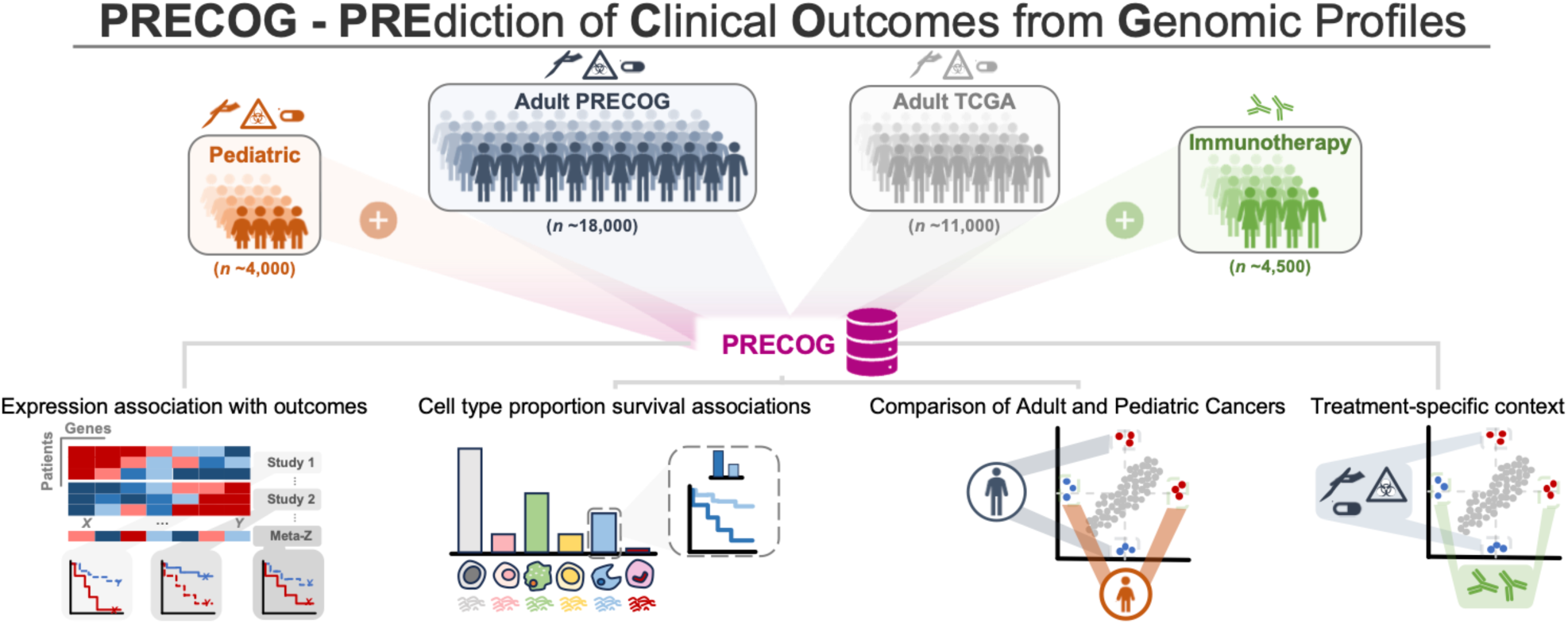

## INTRODUCTION

We previously described the prognostic landscape of genes and infiltrating immune cell types across adult and pediatric cancers and developed an associated freely available resource called PRECOG (PREdiction of Clinical Outcomes from Genomics)^1,2^. PRECOG curated 166 gene expression datasets comprising ∼18,000 tumor samples for which patient outcomes such as overall survival are known. It also includes ∼11,000 samples from TCGA (The Cancer Genome Atlas). The PRECOG resource provides freely downloadable data and annotations, along with precomputed associations between gene expression and outcome, and the ability to visualize these associations through Kaplan-Meier plots.

Although the influence of the immune system on patient responses and survival across cancers and treatment modalities is now well recognized, at the time this was one of the first high-level views of just how pervasive this influence is. In the decade since these resources have been publicly available, immune checkpoint inhibitor (ICI) therapy has revolutionized the treatment of many cancers including, remarkably, in the context of advanced metastatic disease^3^. However, robust prognostic biomarkers that identify which patients will effectively respond to ICI therapies remain elusive for many cancers. For example, tumor mutational burden and parameters such as number of PD1 positive cells are often used, but have limited predictive accuracy^4,5^. Several resources have consolidated gene expression and clinical outcomes for ICI-treated cohorts into databases for data download, exploratory analysis, and hypothesis generation^6–13^. These resources represent useful platforms from which to glean insights into broad and specific predictors of ICI response. However, due to differences across them in areas such as study inclusion criteria (e.g. open source vs. controlled data access), statistical approaches to identify biomarkers of response (e.g. differential expression vs regression), and level of subgrouping across clinical annotations (e.g. considering non small cell lung cancer as a single entity versus separate analysis of adenocarcinoma and squamous cell carcinoma), no single resource is necessarily the most comprehensive or useful compared to the others.

Here we extend PRECOG to the immunotherapy era by performing a comprehensive literature review for studies reporting ICI-treated cancers with transcriptomic profiles and clinical outcomes publicly available for analysis. This search identified numerous studies meeting the previously described criteria. We focused on publicly available data, rather than ones which required data access requests that preclude dissemination - which represents a barrier to open science and reproducibility. Expression associations with ICI response for these reclusive studies have been recently described^11,14^. We also integrate pediatric cancers into PRECOG, facilitating identification of expression biomarkers, and comparison with adult malignancies (Fig 1).

**Figure 1.**
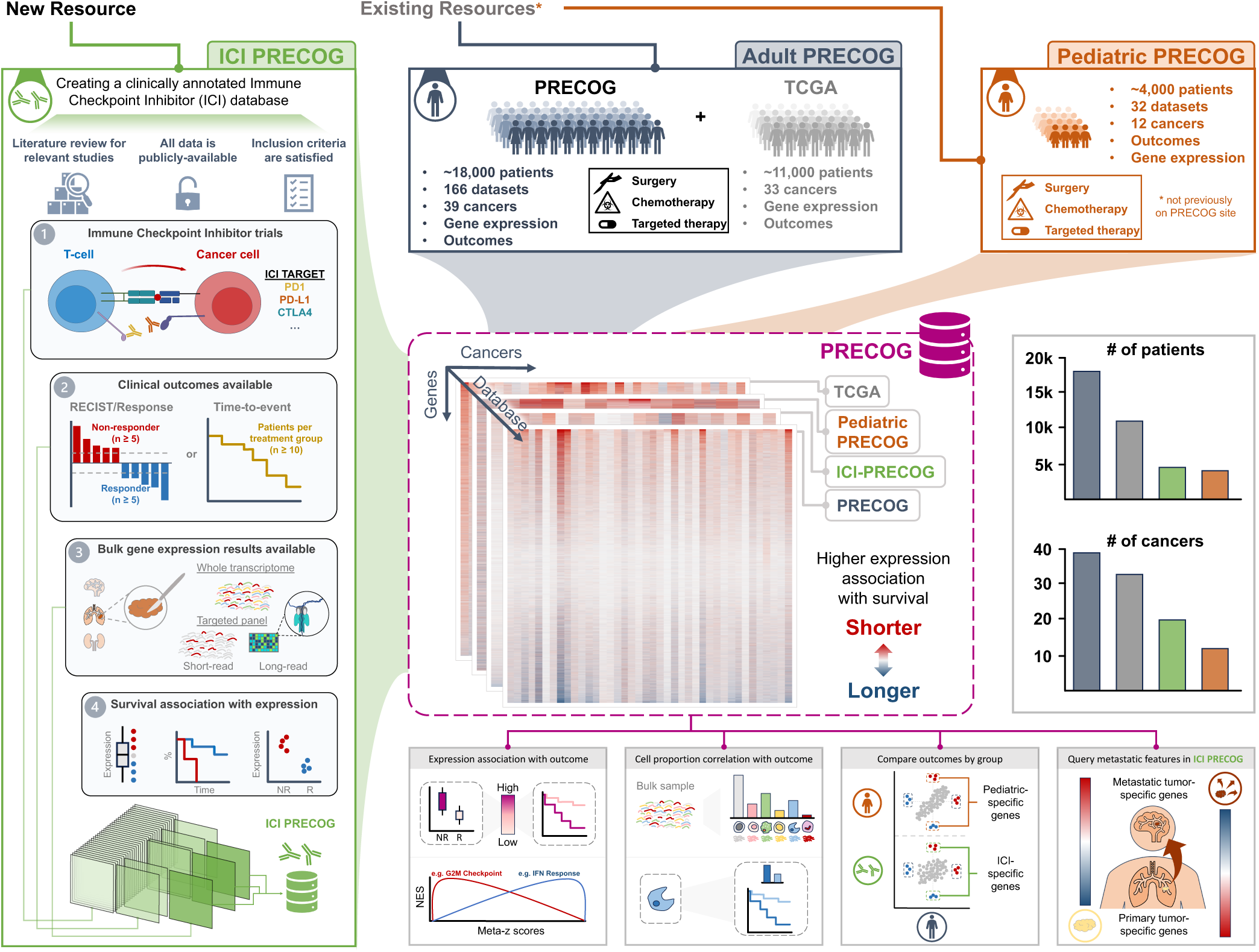
PRECOG update: Integration of existing and novel resources to the site. The PRECOG website previously contained only adult datasets of survival and expression from the original PRECOG publication and TCGA. Our augmentation of the PRECOG site now includes expression and survival associations for ∼4,000 pediatric samples across 12 cancers, in addition to ∼4,500 ICI-treated adult samples across 20 cancers. Precomputed gene expression and cell type abundance correlations with outcomes for individual studies, as well as meta-z scores across dataset, allow comparison between cancers, adult and pediatric malignancies, and treatment modalities.

## MATERIAL AND METHODS

### Dataset curation

#### PRECOG Update

At the time of publication, PRECOG^1^ included only one dataset for pancreatic cancer (PDAC: GSE215011^15^; n = 102) compared with at least two for most other cancers. To improve the signal-to-noise and robustness of associations in PDAC, we added six new PDAC studies with gene expression and clinical outcomes publicly available through GEO: GSE224564^16^ (n = 175), GSE79668^17^ (n = 51), GSE183795^18^ (n = 134), GSE205154^19^ (n = 218), GSE62452^20^ (n = 65), and GSE71729^21^ (n = 125). In combination, these increase the number of PDAC samples from 102 to 870. Gene-level z-scores were generated for each study as previously described^1^ and new PDAC meta-z scores were calculated for all genes across the seven datasets (metaZfunction in package WGCNA v1.73). Curation and analysis of Pediatric PRECOG has been previously described^2^. We also updated annotations for head and neck cancer. Previously, these had not been separated into distinct disease subtypes in PRECOG. We now more rigorously separated entities such as oral squamous carcinoma and hypopharyngeal carcinoma, as described previously^22^. All clinical metadata related to survival were updated including: demographic information (age, sex etc); clinicopathological variables such as tumor subsite, HPV status; and data pertaining to HNC-related risk habits including smoking and alcohol consumption status and intensity measures.

#### Immune Checkpoint Inhibitor (ICI) PRECOG

Studies are included if patients were (1) treated with an immune checkpoint inhibitor, (2) profiled by bulk gene expression, (3) had clinical outcome data such as time-to-event or clinical response available, and (4) all data were publicly accessible. Studies were included regardless of the number of genes measured (i.e. targeted panels vs unbiased assays such as RNA-seq). If studies contained clinically distinct subtypes/histologies (e.g. adenocarcinoma vs squamous in NSCLC), treatment arms, timepoints, sample site (e.g. primary vs metastatic) etc., the study was subset into distinct datasets which were analyzed separately. For studies with time-to-event outcomes data (e.g. overall or progression-free survival, OS/PFS), we excluded datasets with less than ten patients per dataset.

Response Evaluation Criteria in Solid Tumors (RECIST) categories were collapsed to either Responder (complete response CR, partial response PR) or Non-Responder (stable disease SD, progressive disease PD). For studies with RECIST or responder status available, we excluded datasets with less than five patients per response group.

### Clinical metadata

Clinical annotations for all samples were either retrieved from the Gene Expression Omnibus (GEO; package GEOquery v2.70.0), from the study supplementary material, or from publication GitHub repositories^23–26^. All studies were manually cleaned to maintain a consistent nomenclature and labeling across studies as follows: For all studies, we ensure each sample is annotated with respect to primary vs metastatic status, treatment exposure, ICI target, sample timepoint, and cancer type/subtype. In cases of ambiguous, inconsistent, or incomplete sample labeling as provided by the authors, annotations were assigned manually by either extracting the relevant metadata from the manuscript or by recapitulating analyses to assign labels.

### Expression pre-processing and Z-score calculation

#### Pre-processing of expression matrices

For each study, reported gene IDs were updated to their currently approved HGNC HUGO IDs (package biomaRt v2.58.2). Any duplicate gene IDs were combined by taking their average expression level. Next, we (1) ensure the expression matrix is in log2 space, (2) remove all genes with no variance (SD < 0.00001), (3) remove genes/samples with ≥80% missing values, (4) quantile normalize, (5) standardize (scale each gene to have mean zero and unit variance across samples), and (6) impute missing values (package impute v1.76.0). Further details and motivation are provided in the original PRECOG study.

#### Calculating z-scores for gene expression association with outcomes

For studies with sufficient time-to-event outcomes data (n ≥ 10 PFS/OS), univariate Cox proportional hazards regression was used to calculate gene-level z-scores for expression associations with outcomes (package Survival v3.5-8). For studies with responder/non-responder status (n ≥ 5 per group), we used univariate logistic regression to calculate gene-level z-scores for expression associations with responses (package Stats v4.3.1). Positive z-scores indicate that increased gene expression is associated with worse outcomes (i.e. shorter survival time or time to progression, or non-response to therapy). Z-scores were calculated for all outcomes labels when available; for datasets with more than one outcome measure (i.e. OS and PFS, OS and Response), we report all dataset z-scores for the different outcomes measures, prioritizing z-scores for analysis in the following order: OS>PFS>Response.

#### Comparison of Cox proportional hazards z-scores with logistic regression z-scores

PRECOG and Pediatric PRECOG z-scores were all generated from PFS or OS data using Cox proportional hazards regression. ICI PRECOG contains studies with a mixture of OS/PFS, RECIST, and/or responder/non-responder clinical annotations. In order to justify combining z-scores calculated from Cox proportional hazards regression and those calculated using logistic regression, we compared z-scores (for both gene and cell type proportion) from both methods in studies with both time-to-event and responder/non-responder status (n = 25 datasets).

### Cell type abundances and associations with outcome through CIBERSORTx deconvolution

#### Estimating cell type proportions and z-scores

CIBERSORTx^27^ was run on all datasets using default parameters, with the LM22 signature matrix which can enumerate the proportions of 22 different immune cell subtypes in a bulk expression profile. For each dataset with sufficient time-to-event or responder/non-responder status, univariate Cox or logistic regression (respectively) were used to calculate z-scores for each CIBERSORTx cell type fraction and outcomes, analogously to the analysis performed at the level of individual genes. Positive z-scores indicate that increased cell type proportion is associated with worse outcomes/response.

## RESULTS

### Updated PDAC meta z-scores in PRECOG

We added six additional treatment naive primary PDAC datasets (n = 768, Fig 2A) to the single (n=102) study in the original PRECOG. This increased the gene-level meta z-score range from −5.1–4.6 to −8.5–9.3. Gene z-scores between the new and old PDAC meta-z were correlated across 20,610 genes (R = 0.51, p<2.2e-16; Fig 2B). Hallmark genesets^28^ that were statistically associated with outcome (i.e. enriched for individual genes associated with outcome) in the original PRECOG PDAC data all maintained the directionality of association but with increased significance in the updated PRECOG (Fig 2C). Importantly, 26 hallmark sets (out of 50 total tested) that previously did not reach statistical significance were significant (adjusted p<0.05) in the updated PDAC compendium. These included “IFNa Response” and “TNFA Signaling via NFKB” as being associated with worse outcomes in PDAC (Fig 2C), consistent with known biology of PDAC disease^29–31^. There were no examples where genesets lost significance through augmentation of PDAC studies. This illustrates that the updated PRECOG provides an improved resource for researchers to identify clinically-relevant biomarkers and investigate biological processes underlying outcomes.

**Figure 2.**
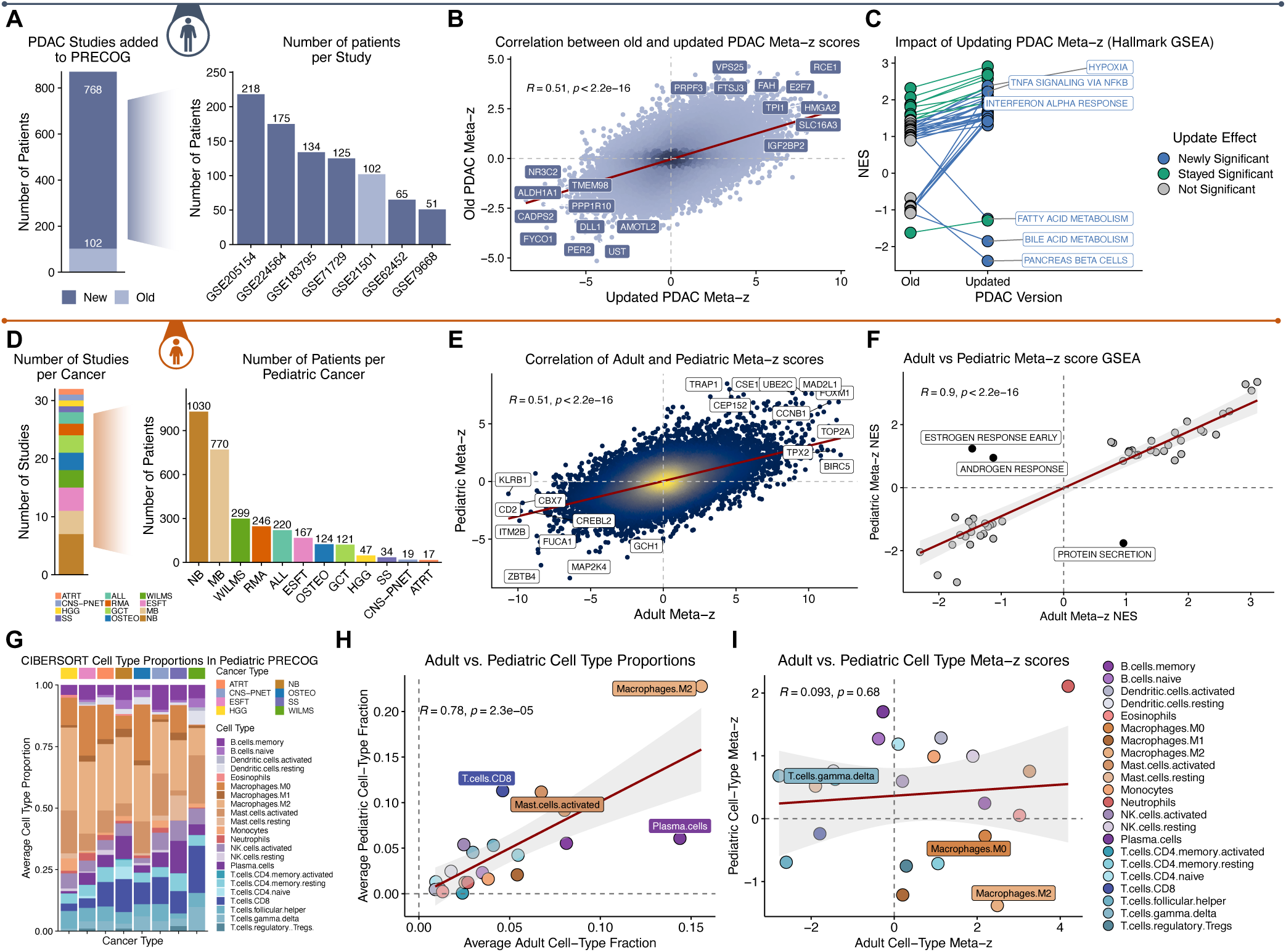
Impact of augmenting adult PRECOG, and comparison to pediatric PRECOG. A) Additional PDAC datasets increase coverage from 102 to 870 patients. B) Old vs updated gene-wise meta-Z scores for PDAC are correlated and show improved significance for many genes which achieve higher absolute Z. C) Geneset-level analysis of old and augmented PDAC data boosts significance of PDAC-relevant pathways and processes associated with survival. D) Summary of number of pediatric cancer studies and cohort sizes. Cancer abbreviations: acute lymphatic leukemia (ALL), atypical teratoid rhabdoid tumor (ATRT), Ewing’s sarcoma family of tumors (ESFT), germ cell tumors (GCT), high grade gliomas (HGG), medulloblastomas (MB), neuroblastoma (NB), osteosarcoma (OSTEO), central nervous system primitive neuroectodermal tumors (CNS-PNETs), rhabdomyosarcoma (RMS), synovial sarcomas (SS) and Wilms tumors (WILMS). E) Comparison between gene-level z-score associations between adult and pediatric cancers shows significant correlation (R=0.51, p<2.2×10-16). F) Hallmarks gene set associations with outcome are highly correlated between adult and pediatric cancers with the exception of protein secretion and adult tissue related development (androgen and estrogen signaling). G) Summary of immune cell type fractions across pediatric cancer types. H) Average cell type fractions across adult cancers and across pediatric cancer are well correlated except M2 macrophages and CD8 T cells (higher in pediatric) and plasma cells (higher in adult). I) Associations between cell type fractions and outcome are discordant between adult and pediatric cancer except neutrophils which are generally unfavorable in both.

### Pediatric PRECOG

#### Pediatric PRECOG overview

Pediatric cancers remain understudied^32^. We previously generated a comprehensive resource of gene expression, estimated cell type proportions, and survival associations across 32 studies of 4,068 patients for 12 pediatric cancers (Pediatric PRECOG; Fig 2D)^2^. However, Pediatric PRECOG was not previously available on https://precog.stanford.edu. To make this resource usable to the broader community, we added all datasets and annotations, study/gene-level survival z-scores, CIBERSORT cell type proportions, and cell type proportion z-scores^2^ to the site. Comparison of pediatric and adult gene-level meta-z scores showed significant correlation (R = 0.51, p<2.2e-16; Fig 2E). Interestingly, GSEA enrichment scores for Hallmark pathways between adult and pediatric meta-z scores were highly correlated (R = 0.9, p<2.2e-16; Fig 2F), with the exception of “Estrogen/Androgen Response” and “Protein Secretion”. These make sense as they are fundamental differences in biology and development between pediatric and adult cancers, with estrogen and androgen receptor signaling playing key roles driving cancer-relevant processes such as proliferation in adult tissues.

We estimated the proportions of 22 leukocyte cell types^33^ across Pediatric PRECOG datasets as previously described^2^ (Fig 1G). Comparing adult and pediatric cancer, the average proportions of most cell types were highly correlated (R = 0.78, p = 2.3×10-5; Fig 2H). However, the association of cell type proportion with outcomes was uncorrelated, with neutrophils being the only cell type with a consistent, unfavorable, association with outcome between adult and pediatric cancers (Fig 2I). The neutrophil signal potentially clarifies their increasingly controversial impact of inflammation in cancer^34,35^. This provides one example highlighting the utility of Pediatric PRECOG and provides a rich resource for uncovering developmentally conserved and distinct biomarkers for further research.

### ICI PRECOG

#### ICI PRECOG cohort description

Our literature review for immunotherapy studies with publicly-available gene expression and clinical outcomes data for treatment naive patients identified 4,045 patients across 51 studies and 20 cancers, of which 851 samples were lymph node or distant metastases (Fig 3A-C)^36–87^. We subset “studies” into “datasets” if there were at least 10 samples with the same cancer subtype (e.g. adenocarcinoma vs squamous), ICI target, primary/metastatic annotation, and treatment time point (e.g. naive vs post therapy). This defined 69 datasets for which we calculated outcome-associated gene z-scores (49 primary naive, 20 metastatic naive). To define prognostic associations for distinct clinical categories, we collapsed ICI datasets based on shared cancer type, ICI target, primary/metastatic status, and treatment exposure time point (e.g. pre/post therapy) and calculated meta-z scores for each category (Fig 3D). For datasets for which both OS/PFS and binary response were available, we compared gene-level z-scores and observed that they were well correlated (Fig 3. D; R=0.73, p=0.0004).

**Figure 3.**
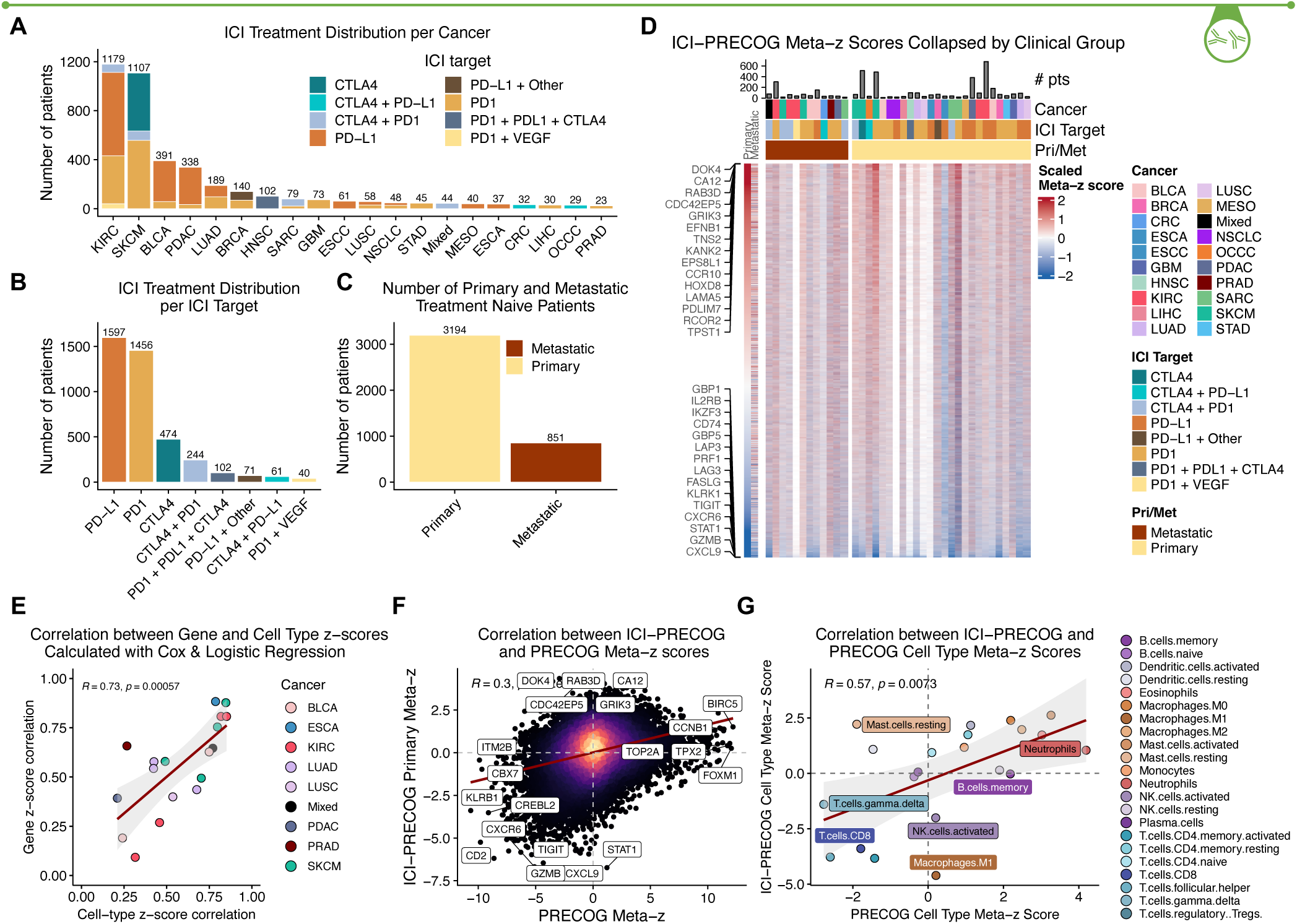
Development and high level analysis of ICI PRECOG. A) Summary of the number of patients per cancer type, together with specific ICI treatment. B) Distribution of ICI treatments across ICI PRECOG datasets. C) ICI PRECOG distribution of primary versus metastatic samples. D) Top-level heatmap of ICI PRECOG gene expression associations with outcome. Red indicates that a gene (row) is associated with worse survival (Cox regression) or binary lack-of-response (logistic regression) in a specific dataset (column). Cancer types correspond to TCGA designations. E) Comparison between gene-level and cell-type-level z-scores for datasets where both time-to-event and binary response are available. F) Gene-level z-scores are modestly correlated between Adult PRECOG (non-ICI therapy) and ICI PRECOG (R=0.3, p<2,2×10-16), with most agreement occurring in favorable genes (bottom left quadrant) which primarily reflect immune activity. G) Concordance between the association of specific immune cell types with outcome in Adult PRECOG vs ICI PRECOG shows significant correlation except for mast cells and M1 macrophages.

#### ICI PRECOG compared to PRECOG

When gene expression levels were associated with ICI response, many known “T cell exhaustion” genes such as *TIGIT* and *LAG3* emerged as favorable prognostic markers; while the single most unfavorable marker for ICI response was *DOK4* which encodes for an adapter protein that has recently been implicated in negative regulation of T-cell activation^88^. Gene-level meta-z scores showed modest correlation between ICI PRECOG and PRECOG (R = 0.3, p < 2.2e-19; Fig 1F). Immune related genes were often favorably prognostic in both ICI and non-ICI contexts; whereas genes associated with poor survival differed between them. Hence, ICI PRECOG retrieves both familiar and also less-known markers of immunotherapy response, including ones that are shared or distinct from those that are prognostic for older therapies. This illustrates the potential for discovery of new markers and mechanisms underlying ICI response. We also estimated proportions of 22 different immune cell types for each sample in ICI PRECOG and computed z-scores for their association with ICI response or survival. These were broadly correlated with similar measures in the original PRECOG (Fig 3G), with neutrophils being unfavorable in both, and CD8 T cells and γδT cells favorable in both. However, there were strong differences for some cell types, such as M1 macrophages and mast cells, whose presence was unfavorable in the context of ICI therapy, but otherwise favorable.

## DISCUSSION

Here, we integrated and augmented our PRECOG database of adult^1^ and pediatric cancers^2^; and added a large curated, processed, and pre-analyzed compendium of immunotherapy response datasets of over 4,000 patients. All data and annotations are freely available for download from the PRECOG website at https://precog.stanford.edu. This resource also provides capability for basic analyses such as comparing gene z-score association with outcomes between cancer types and individual datasets. The updated PRECOG represents a significant centralization of gene expression and cell type abundance associations with outcomes across pediatric and adult cancers with immunotherapy treatment. While there are similar resources for ICI treated cohorts^6–13^, PRECOG is the most comprehensive in terms of treatment types, data availability, consistent and curated sample annotations, and the ability to integrate with non-ICI cohorts, as well as comparing and contrasting to pediatric cancer. The vignettes we present here illustrate the potential for discovery of new biomarkers related to ICI response. However, we also anticipate that more in-depth analysis, paired with comparison to older therapies, will be informative about mechanisms of response and resistance development.

## DATA AVAILABILITY

Only publicly available datasets were used in this study. All clinical and expression data used, as well as dataset level CIBERSORT(x) cell-type fractions and gene/cell-type z-scores generated in this study can be downloaded at https://precog.stanford.edu.

## AUTHOR CONTRIBUTIONS

Brooks Benard: Conceptualization, Data curation, Formal analysis, Methodology, Project administration, Software, Supervision, Visualization, Writing—original draft. Chinmay Lalgudi: Resource, Software, Visualization, Writing—review & editing. Ilayda Ilerten: Formal analysis, Visualization. Ruohan Wang: Formal analysis. Andrew Gentles: Conceptualization, Formal analysis, Funding acquisition, Project administration, Methodology, Supervision, Writing—review & editing.

## ACKNOWLEDGEMENTS

We would like to thank all the authors of the included studies for making their data publicly available and useful to the scientific community.

## REFERENCES

(1) Gentles, A. J.; Newman, A. M.; Liu, C. L.; Bratman, S. V.; Feng, W.; Kim, D.; Nair, V. S.; Xu, Y.; Khuong, A.; Hoang, C. D.; Diehn, M.; West, R. B.; Plevritis, S. K.; Alizadeh, A. A. The Prognostic Landscape of Genes and Infiltrating Immune Cells across Human Cancers. Nat. Med. 2015, 21 (8), 938–945. 10.1038/nm.3909.

(2) Stahl, D.; Knoll, R.; Gentles, A. J.; Vokuhl, C.; Buness, A.; Gütgemann, I. Prognostic Gene Expression, Stemness and Immune Microenvironment in Pediatric Tumors. Cancers 2021, 13 (4), 854. 10.3390/cancers13040854.

(3) Ganesh, K.; Massagué, J. Targeting Metastatic Cancer. Nat. Med. 2021, 27 (1), 34–44. 10.1038/s41591-020-01195-4.

(4) Davis, A. A.; Patel, V. G. The Role of PD-L1 Expression as a Predictive Biomarker: An Analysis of All US Food and Drug Administration (FDA) Approvals of Immune Checkpoint Inhibitors. J. Immunother. Cancer 2019, 7 (1). 10.1186/s40425-019-0768-9.

(5) Li, H.; van der Merwe, P. A.; Sivakumar, S. Biomarkers of Response to PD-1 Pathway Blockade. Br. J. Cancer 2022, 126 (12), 1663–1675. 10.1038/s41416-022-01743-4.

(6) Chen, Z.; Luo, Z.; Zhang, D.; Li, H.; Liu, X.; Zhu, K.; Zhang, H.; Wang, Z.; Zhou, P.; Ren, J.; Zhao, A.; Zuo, Z. TIGER: A Web Portal of Tumor Immunotherapy Gene Expression Resource. Genomics Proteomics Bioinformatics 2023, 21 (2), 337–348. 10.1016/j.gpb.2022.08.004.

(7) Eddy, J. A.; Thorsson, V.; Lamb, A. E.; Gibbs, D. L.; Heimann, C.; Yu, J. X.; Chung, V.; Chae, Y.; Dang, K.; Vincent, B. G.; Shmulevich, I.; Guinney, J. CRI iAtlas: An Interactive Portal for Immuno-Oncology Research. F1000Research August 24, 2020. 10.12688/f1000research.25141.1.

(8) Liu, Y.; Zhou, Y.; Hu, X.; Le-Ge, W.; Wang, H.; Jiang, T.; Li, J.; Hu, Y.; Wang, Y. DIRMC: A Database of Immunotherapy-Related Molecular Characteristics. Database 2024, 2024, baae032. 10.1093/database/baae032.

(9) Yang, J.; Liu, Q.; Shyr, Y. A Large-Scale Meta-Analysis Reveals Positive Feedback between Macrophages and T Cells That Sensitizes Tumors to Immunotherapy. Cancer Res. 2024, 84 (4), 626–638. 10.1158/0008-5472.CAN-23-2006.

(10) Yang, M.; Miao, Y.-R.; Xie, G.-Y.; Luo, M.; Hu, H.; Kwok, H. F.; Feng, J.; Guo, A.-Y. ICBatlas: A Comprehensive Resource for Depicting Immune Checkpoint Blockade Therapy Characteristics from Transcriptome Profiles. Cancer Immunol. Res. 2022, 10 (11), 1398–1406. 10.1158/2326-6066.CIR-22-0249.

(11) Gong, L.; Luo, J.; Yang, E.; Ru, B.; Qi, Z.; Yang, Y.; Rani, A.; Purohit, A.; Zhang, Y.; Guan, G.; Paul, R.; Vu, T.; Chen, Z.; Ji, R.; Day, C.-P.; Wu, C.; Merlino, G.; Fitzgerald, D.; Altan-Bonnet, G.; Aldape, K.; Wu, J.; Guan, X.; Jiang, P. Cancer Immunology Data Engine Reveals Secreted AOAH as a Potential Immunotherapy. Cell 2025, 0 (0). 10.1016/j.cell.2025.07.004.

(12) Zhang, Y.; Yao, Y.; Chen, P.; Liu, Y.; Zhang, H.; Liu, H.; Liu, Y.; Xu, H.; Tian, X.; Wang, Z.; Chu, P.; Zhao, D.; Liu, H.; Zhang, C.; Chen, S.; Zhao, Y.; Liu, C.; Yang, Y. Checkpoint Therapeutic Target Database (CKTTD): The First Comprehensive Database for Checkpoint Targets and Their Modulators in Cancer Immunotherapy. J. Immunother. Cancer 2020, 8 (2). 10.1136/jitc-2020-001247.

(13) Zeng, Z.; Wong, C. J.; Yang, L.; Ouardaoui, N.; Li, D.; Zhang, W.; Gu, S.; Zhang, Y.; Liu, Y.; Wang, X.; Fu, J.; Zhou, L.; Zhang, B.; Kim, S.; Yates, K. B.; Brown, M.; Freeman, G. J.; Uppaluri, R.; Manguso, R.; Liu, X. S. TISMO: Syngeneic Mouse Tumor Database to Model Tumor Immunity and Immunotherapy Response. Nucleic Acids Res. 2022, 50 (D1), D1391–D1397. 10.1093/nar/gkab804.

(14) Lee, J. S.; Nair, N. U.; Dinstag, G.; Chapman, L.; Chung, Y.; Wang, K.; Sinha, S.; Cha, H.; Kim, D.; Schperberg, A. V.; Srinivasan, A.; Lazar, V.; Rubin, E.; Hwang, S.; Berger, R.; Beker, T.; Ronai, Z.; Hannenhalli, S.; Gilbert, M. R.; Kurzrock, R.; Lee, S.-H.; Aldape, K.; Ruppin, E. Synthetic Lethality-Mediated Precision Oncology via the Tumor Transcriptome. Cell 2021, 184 (9), 2487–2502.e13. 10.1016/j.cell.2021.03.030.

(15) Liu, C.; Zhou, C.; Xia, W.; Zhou, Y.; Qiu, Y.; Weng, J.; Zhou, Q.; Chen, W.; Wang, Y.-N.; Lee, H.-H.; Wang, S.-C.; Kuang, M.; Yu, D.; Ren, N.; Hung, M.-C. Targeting ALK Averts Ribonuclease 1-Induced Immunosuppression and Enhances Antitumor Immunity in Hepatocellular Carcinoma. Nat. Commun. 2024, 15 (1), 1009. 10.1038/s41467-024-45215-0.

(16) Ohara, Y.; Tang, W.; Liu, H.; Yang, S.; Dorsey, T. H.; Cawley, H.; Moreno, P.; Chari, R.; Guest, M. R.; Azizian, A.; Gaedcke, J.; Ghadimi, M.; Hanna, N.; Ambs, S.; Hussain, S. P. SERPINB3-MYC Axis Induces the Basal-like/Squamous Subtype and Enhances Disease Progression in Pancreatic Cancer. Cell Rep. 2023, 42 (12), 113434. 10.1016/j.celrep.2023.113434.

(17) Kirby, M. K.; Ramaker, R. C.; Gertz, J.; Davis, N. S.; Johnston, B. E.; Oliver, P. G.; Sexton, K. C.; Greeno, E. W.; Christein, J. D.; Heslin, M. J.; Posey, J. A.; Grizzle, W. E.; Vickers, S. M.; Buchsbaum, D. J.; Cooper, S. J.; Myers, R. M. RNA Sequencing of Pancreatic Adenocarcinoma Tumors Yields Novel Expression Patterns Associated with Long-Term Survival and Reveals a Role for ANGPTL4. Mol. Oncol. 2016, 10 (8), 1169–1182. 10.1016/j.molonc.2016.05.004.

(18) Yang, S.; Tang, W.; Azizian, A.; Gaedcke, J.; Ströbel, P.; Wang, L.; Cawley, H.; Ohara, Y.; Valenzuela, P.; Zhang, L.; Lal, T.; Sinha, S.; Rupin, E.; Hanna, N.; Ghadimi, B. M.; Hussain, S. P. Dysregulation of HNF1B/Clusterin Axis Enhances Disease Progression in a Highly Aggressive Subset of Pancreatic Cancer Patients. Carcinogenesis 2022, 43 (12), 1198–1210. 10.1093/carcin/bgac092.

(19) Link, J. M.; Eng, J. R.; Pelz, C.; MacPherson-Hawthorne, K.; Worth, P. J.; Sivagnanam, S.; Keith, D. J.; Owen, S.; Langer, E. M.; Grossblatt-Wait, A.; Salgado-Garza, G.; Creason, A. L.; Protzek, S.; Egger, J.; Holly, H.; Heskett, M. B.; Chin, K.; Kirchberger, N.; Betre, K.; Bucher, E.; Kilburn, D.; Hu, Z.; Munks, M. W.; English, I. A.; Tsuda, M.; Goecks, J.; Demir, E.; Adey, A. C.; Kardosh, A.; Lopez, C. D.; Sheppard, B. C.; Guimaraes, A.; Brinkerhoff, B.; Morgan, T. K.; Mills, G. B.; Coussens, L. M.; Brody, J. R.; Sears, R. C. Ongoing Replication Stress Tolerance and Clonal T Cell Responses Distinguish Liver and Lung Recurrence and Outcomes in Pancreatic Cancer. Nat. Cancer 2025, 6 (1), 123–144. 10.1038/s43018-024-00881-3.

(20) Yang, S.; He, P.; Wang, J.; Schetter, A.; Tang, W.; Funamizu, N.; Yanaga, K.; Uwagawa, T.; Satoskar, A. R.; Gaedcke, J.; Bernhardt, M.; Ghadimi, B. M.; Gaida, M. M.; Bergmann, F.; Werner, J.; Ried, T.; Hanna, N.; Alexander, H. R.; Hussain, S. P. A Novel MIF Signaling Pathway Drives the Malignant Character of Pancreatic Cancer by Targeting NR3C2. Cancer Res. 2016, 76 (13), 3838–3850. 10.1158/0008-5472.CAN-15-2841.

(21) Moffitt, R. A.; Marayati, R.; Flate, E. L.; Volmar, K. E.; Loeza, S. G. H.; Hoadley, K. A.; Rashid, N. U.; Williams, L. A.; Eaton, S. C.; Chung, A. H.; Smyla, J. K.; Anderson, J. M.; Kim, H. J.; Bentrem, D. J.; Talamonti, M. S.; Iacobuzio-Donahue, C. A.; Hollingsworth, M. A.; Yeh, J. J. Virtual Microdissection Identifies Distinct Tumor- and Stroma-Specific Subtypes of Pancreatic Ductal Adenocarcinoma. Nat. Genet. 2015, 47 (10), 1168–1178. 10.1038/ng.3398.

(22) Brennan, K.; Espín-Pérez, A.; Chang, S.; Bedi, N.; Saumyaa, S.; Shin, J. H.; Plevritis, S. K.; Gevaert, O.; Sunwoo, J. B.; Gentles, A. J. Loss of P53-DREAM-Mediated Repression of Cell Cycle Genes as a Driver of Lymph Node Metastasis in Head and Neck Cancer. Genome Med. 2023, 15 (1), 98. 10.1186/s13073-023-01236-w.

(23) Campbell, K. M.; Amouzgar, M.; Pfeiffer, S. M.; Howes, T. R.; Medina, E.; Travers, M.; Steiner, G.; Weber, J. S.; Wolchok, J. D.; Larkin, J.; Hodi, F. S.; Boffo, S.; Salvador, L.; Tenney, D.; Tang, T.; Thompson, M. A.; Spencer, C. N.; Wells, D. K.; Ribas, A. Prior Anti-CTLA-4 Therapy Impacts Molecular Characteristics Associated with Anti-PD-1 Response in Advanced Melanoma. Cancer Cell 2023, 41 (4), 791–806.e4. 10.1016/j.ccell.2023.03.010.

(24) Padrón, L. J.; Maurer, D. M.; O’Hara, M. H.; O’Reilly, E. M.; Wolff, R. A.; Wainberg, Z. A.; Ko, A. H.; Fisher, G.; Rahma, O.; Lyman, J. P.; Cabanski, C. R.; Yu, J. X.; Pfeiffer, S. M.; Spasic, M.; Xu, J.; Gherardini, P. F.; Karakunnel, J.; Mick, R.; Alanio, C.; Byrne, K. T.; Hollmann, T. J.; Moore, J. S.; Jones, D. D.; Tognetti, M.; Chen, R. O.; Yang, X.; Salvador, L.; Wherry, E. J.; Dugan, U.; O’Donnell-Tormey, J.; Butterfield, L. H.; Hubbard-Lucey, V. M.; Ibrahim, R.; Fairchild, J.; Bucktrout, S.; LaVallee, T. M.; Vonderheide, R. H. Sotigalimab and/or Nivolumab with Chemotherapy in First-Line Metastatic Pancreatic Cancer: Clinical and Immunologic Analyses from the Randomized Phase 2 PRINCE Trial. Nat. Med. 2022, 28 (6), 1167–1177. 10.1038/s41591-022-01829-9.

(25) Galsky, M. D.; Autio, K. A.; Cabanski, C. R.; Wentzel, K.; Graff, J. N.; Friedlander, T. W.; Howes, T. R.; Shotts, K. M.; Densmore, J.; Spasic, M.; Da Silva, D. M.; Chen, R. O.; Lata, J.; Skolnik, J.; Keler, T.; Yellin, M. J.; LaVallee, T. M.; Fairchild, J.; Boffo, S.; O’Donnell-Tormey, J.; Dugan, U.; Bhardwaj, N.; Subudhi, S. K.; Fong, L. Clinical and Translational Results from PORTER, a Multicohort Phase I Platform Trial of Combination Immunotherapy in Metastatic Castration-Resistant Prostate Cancer. Clin. Cancer Res. 2025, 31 (8), 1463–1475. 10.1158/1078-0432.CCR-24-3693.

(26) Cui, C.; Xu, C.; Yang, W.; Chi, Z.; Sheng, X.; Si, L.; Xie, Y.; Yu, J.; Wang, S.; Yu, R.; Guo, J.; Kong, Y. Ratio of the Interferon-γ Signature to the Immunosuppression Signature Predicts Anti-PD-1 Therapy Response in Melanoma. Npj Genomic Med. 2021, 6 (1), 7. 10.1038/s41525-021-00169-w.

(27) Newman, A. M.; Steen, C. B.; Liu, C. L.; Gentles, A. J.; Chaudhuri, A. A.; Scherer, F.; Khodadoust, M. S.; Esfahani, M. S.; Luca, B. A.; Steiner, D.; Diehn, M.; Alizadeh, A. A. Determining Cell Type Abundance and Expression from Bulk Tissues with Digital Cytometry. Nat. Biotechnol. 2019, 37 (7), 773–782. 10.1038/s41587-019-0114-2.

(28) Liberzon, A.; Birger, C.; Thorvaldsdóttir, H.; Ghandi, M.; Mesirov, J. P.; Tamayo, P. The Molecular Signatures Database Hallmark Gene Set Collection. Cell Syst. 2015, 1 (6), 417–425. 10.1016/j.cels.2015.12.004.

(29) Adjuto-Saccone, M.; Soubeyran, P.; Garcia, J.; Audebert, S.; Camoin, L.; Rubis, M.; Roques, J.; Binétruy, B.; Iovanna, J. L.; Tournaire, R. TNF-α Induces Endothelial–Mesenchymal Transition Promoting Stromal Development of Pancreatic Adenocarcinoma. Cell Death Dis. 2021, 12 (7), 649. 10.1038/s41419-021-03920-4.

(30) Alam, M. S.; Gaida, M. M.; Witzel, H. R.; Otsuka, S.; Abbasi, A.; Guerin, T.; Abdelmaksoud, A.; Wong, N.; Cam, M. C.; Kozlov, S.; Ashwell, J. D. TNFR1 Signaling Promotes Pancreatic Tumor Growth by Limiting Dendritic Cell Number and Function. Cell Rep. Med. 2024, 5 (9), 101696. 10.1016/j.xcrm.2024.101696.

(31) Espinet, E.; Gu, Z.; Imbusch, C. D.; Giese, N. A.; Büscher, M.; Safavi, M.; Weisenburger, S.; Klein, C.; Vogel, V.; Falcone, M.; Insua-Rodríguez, J.; Reitberger, M.; Thiel, V.; Kossi, S. O.; Muckenhuber, A.; Sarai, K.; Lee, A. Y. L.; Backx, E.; Zarei, S.; Gaida, M. M.; Rodríguez-Paredes, M.; Donato, E.; Yen, H.-Y.; Eils, R.; Schlesner, M.; Pfarr, N.; Hackert, T.; Plass, C.; Brors, B.; Steiger, K.; Weichenhan, D.; Arda, H. E.; Rooman, I.; Kopp, J. L.; Strobel, O.; Weichert, W.; Sprick, M. R.; Trumpp, A. Aggressive PDACs Show Hypomethylation of Repetitive Elements and the Execution of an Intrinsic IFN Program Linked to a Ductal Cell of Origin. Cancer Discov. 2021, 11 (3), 638–659. 10.1158/2159-8290.CD-20-1202.

(32) Gallicchio, L.; Mollica, M.; Tesauro, G.; Doose, M.; Guida, J. L.; Maher, M. E.; Tonorezos, E. Rare Cancer Survivorship Research Funding at the National Institutes of Health (NIH), 2017 to 2023. Cancer Causes Control 2025, 36 (6), 587–594. 10.1007/s10552-025-01959-8.

(33) Chen, B.; Khodadoust, M. S.; Liu, C. L.; Newman, A. M.; Alizadeh, A. A. Profiling Tumor Infiltrating Immune Cells with CIBERSORT. In Cancer Systems Biology: Methods and Protocols; von Stechow, L., Ed.; Springer: New York, NY, 2018; pp 243–259. 10.1007/978-1-4939-7493-1_12.

(34) Xiong, S.; Dong, L.; Cheng, L. Neutrophils in Cancer Carcinogenesis and Metastasis. J. Hematol. Oncol.J Hematol Oncol 2021, 14 (1), 173. 10.1186/s13045-021-01187-y.

(35) Huang, X.; Nepovimova, E.; Adam, V.; Sivak, L.; Heger, Z.; Valko, M.; Wu, Q.; Kuca, K. Neutrophils in Cancer Immunotherapy: Friends or Foes? Mol. Cancer 2024, 23 (1), 107. 10.1186/s12943-024-02004-z.

(36) Tsimberidou, A. M.; Alayli, F. A.; Okrah, K.; Drakaki, A.; Khalil, D. N.; Kummar, S.; Khan, S. A.; Hodi, F. S.; Oh, D. Y.; Cabanski, C. R.; Gautam, S.; Meier, S. L.; Amouzgar, M.; Pfeiffer, S. M.; Kageyama, R.; Yang, E.; Spasic, M.; Tetzlaff, M. T.; Foo, W. C.; Hollmann, T. J.; Li, Y.; Adamow, M.; Wong, P.; Moore, J. S.; Velichko, S.; Chen, R. O.; Kumar, D.; Bucktrout, S.; Ibrahim, R.; Dugan, U.; Salvador, L.; Hubbard-Lucey, V. M.; O’Donnell-Tormey, J.; Santulli-Marotto, S.; Butterfield, L. H.; Da Silva, D. M.; Fairchild, J.; LaVallee, T. M.; Padrón, L. J.; Sharma, P. Immunologic Signatures of Response and Resistance to Nivolumab with Ipilimumab in Advanced Metastatic Cancer. J. Exp. Med. 2024, 221 (10), e20240152. 10.1084/jem.20240152.

(37) Weber, J. S.; Gibney, G.; Sullivan, R. J.; Sosman, J. A.; Slingluff, C. L.; Lawrence, D. P.; Logan, T. F.; Schuchter, L. M.; Nair, S.; Fecher, L.; Buchbinder, E. I.; Berghorn, E.; Ruisi, M.; Kong, G.; Jiang, J.; Horak, C.; Hodi, F. S. Sequential Administration of Nivolumab and Ipilimumab with a Planned Switch in Patients with Advanced Melanoma (CheckMate 064): An Open-Label, Randomised, Phase 2 Trial. Lancet Oncol. 2016, 17 (7), 943–955. 10.1016/S1470-2045(16)30126-7.

(38) Wolchok, J. D.; Chiarion-Sileni, V.; Gonzalez, R.; Rutkowski, P.; Grob, J.-J.; Cowey, C. L.; Lao, C. D.; Wagstaff, J.; Schadendorf, D.; Ferrucci, P. F.; Smylie, M.; Dummer, R.; Hill, A.; Hogg, D.; Haanen, J.; Carlino, M. S.; Bechter, O.; Maio, M.; Marquez-Rodas, I.; Guidoboni, M.; McArthur, G.; Lebbé, C.; Ascierto, P. A.; Long, G. V.; Cebon, J.; Sosman, J.; Postow, M. A.; Callahan, M. K.; Walker, D.; Rollin, L.; Bhore, R.; Hodi, F. S.; Larkin, J. Overall Survival with Combined Nivolumab and Ipilimumab in Advanced Melanoma. N. Engl. J. Med. 2017, 377 (14), 1345–1356. 10.1056/NEJMoa1709684.

(39) Braun, D. A.; Hou, Y.; Bakouny, Z.; Ficial, M.; Sant’ Angelo, M.; Forman, J.; Ross-Macdonald, P.; Berger, A. C.; Jegede, O. A.; Elagina, L.; Steinharter, J.; Sun, M.; Wind-Rotolo, M.; Pignon, J.-C.; Cherniack, A. D.; Lichtenstein, L.; Neuberg, D.; Catalano, P.; Freeman, G. J.; Sharpe, A. H.; McDermott, D. F.; Van Allen, E. M.; Signoretti, S.; Wu, C. J.; Shukla, S. A.; Choueiri, T. K. Interplay of Somatic Alterations and Immune Infiltration Modulates Response to PD-1 Blockade in Advanced Clear Cell Renal Cell Carcinoma. Nat. Med. 2020, 26 (6), 909–918. 10.1038/s41591-020-0839-y.

(40) Chen, P.-L.; Roh, W.; Reuben, A.; Cooper, Z. A.; Spencer, C. N.; Prieto, P. A.; Miller, J. P.; Bassett, R. L.; Gopalakrishnan, V.; Wani, K.; De Macedo, M. P.; Austin-Breneman, J. L.; Jiang, H.; Chang, Q.; Reddy, S. M.; Chen, W.-S.; Tetzlaff, M. T.; Broaddus, R. J.; Davies, M. A.; Gershenwald, J. E.; Haydu, L.; Lazar, A. J.; Patel, S. P.; Hwu, P.; Hwu, W.-J.; Diab, A.; Glitza, I. C.; Woodman, S. E.; Vence, L. M.; Wistuba, I. I.; Amaria, R. N.; Kwong, L. N.; Prieto, V.; Davis, R. E.; Ma, W.; Overwijk, W. W.; Sharpe, A. H.; Hu, J.; Futreal, P. A.; Blando, J.; Sharma, P.; Allison, J. P.; Chin, L.; Wargo, J. A. Analysis of Immune Signatures in Longitudinal Tumor Samples Yields Insight into Biomarkers of Response and Mechanisms of Resistance to Immune Checkpoint Blockade. Cancer Discov. 2016, 6 (8), 827–837. 10.1158/2159-8290.CD-15-1545.

(41) Dai, Y.; Knisely, A.; Yano, M.; Dang, M.; Hinchcliff, E. M.; Lee, S.; Welp, A.; Chelvanambi, M.; Lastrapes, M.; Liu, H.; Yuan, Z.; Wang, C.; Nie, H.; Jean, S.; Montaner, L. J.; Hou, J.; Patel, A.; Patel, S.; Fellman, B.; Yuan, Y.; Sun, B.; Pandurengan, R. K.; Cuentas, E. R. P.; Celestino, J.; Liu, Y.; Liu, J.; Hillman, R. T.; Westin, S. N.; Sood, A. K.; Soliman, P. T.; Shafer, A.; Meyer, L. A.; Gershenson, D. M.; Vining, D.; Ganeshan, D.; Lu, K.; Wargo, J. A.; Peng, W.; Zhang, R.; Wang, L.; Jazaeri, A. A. PPP2R1A Mutations Portend Improved Survival after Cancer Immunotherapy. Nature 2025, 644 (8076), 537–546. 10.1038/s41586-025-09203-8.

(42) Potential markers of response and resistance to programmed cell death-1 blockade in first-line therapy of cisplatin-inilegible advanced urothelial cancer. - ASCO. https://www.asco.org/abstracts-presentations/ABSTRACT244127 (accessed 2025-08-15).

(43) Huang, A. C.; Orlowski, R. J.; Xu, X.; Mick, R.; George, S. M.; Yan, P. K.; Manne, S.; Kraya, A. A.; Wubbenhorst, B.; Dorfman, L.; D’Andrea, K.; Wenz, B. M.; Liu, S.; Chilukuri, L.; Kozlov, A.; Carberry, M.; Giles, L.; Kier, M. W.; Quagliarello, F.; McGettigan, S.; Kreider, K.; Annamalai, L.; Zhao, Q.; Mogg, R.; Xu, W.; Blumenschein, W. M.; Yearley, J. H.; Linette, G. P.; Amaravadi, R. K.; Schuchter, L. M.; Herati, R. S.; Bengsch, B.; Nathanson, K. L.; Farwell, M. D.; Karakousis, G. C.; Wherry, E. J.; Mitchell, T. C. A Single Dose of Neoadjuvant PD-1 Blockade Predicts Clinical Outcomes in Resectable Melanoma. Nat. Med. 2019, 25 (3), 454–461. 10.1038/s41591-019-0357-y.

(44) Kim, J. Y.; Choi, J. K.; Jung, H. Genome-Wide Methylation Patterns Predict Clinical Benefit of Immunotherapy in Lung Cancer. Clin. Epigenetics 2020, 12 (1), 119. 10.1186/s13148-020-00907-4.

(45) Hwang, S.; Kwon, A.-Y.; Jeong, J.-Y.; Kim, S.; Kang, H.; Park, J.; Kim, J.-H.; Han, O. J.; Lim, S. M.; An, H. J. Immune Gene Signatures for Predicting Durable Clinical Benefit of Anti-PD-1 Immunotherapy in Patients with Non-Small Cell Lung Cancer. Sci. Rep. 2020, 10 (1), 643. 10.1038/s41598-019-57218-9.

(46) Hsu, C.-L.; Ou, D.-L.; Bai, L.-Y.; Chen, C.-W.; Lin, L.; Huang, S.-F.; Cheng, A.-L.; Jeng, Y.-M.; Hsu, C. Exploring Markers of Exhausted CD8 T Cells to Predict Response to Immune Checkpoint Inhibitor Therapy for Hepatocellular Carcinoma. Liver Cancer 2021, 10 (4), 346–359. 10.1159/000515305.

(47) Foy, J.-P.; Karabajakian, A.; Ortiz-Cuaran, S.; Boussageon, M.; Michon, L.; Bouaoud, J.; Fekiri, D.; Robert, M.; Baffert, K.-A.; Hervé, G.; Quilhot, P.; Attignon, V.; Girod, A.; Chaine, A.; Benassarou, M.; Zrounba, P.; Caux, C.; Ghiringhelli, F.; Lantuejoul, S.; Crozes, C.; Brochériou, I.; Pérol, M.; Fayette, J.; Bertolus, C.; Saintigny, P. Immunologically Active Phenotype by Gene Expression Profiling Is Associated with Clinical Benefit from PD-1/PD-L1 Inhibitors in Real-World Head and Neck and Lung Cancer Patients. Eur. J. Cancer 2022, 174, 287–298. 10.1016/j.ejca.2022.06.034.

(48) Foy, J.-P.; Karabajakian, A.; Ortiz-Cuaran, S.; Boussageon, M.; Michon, L.; Bouaoud, J.; Fekiri, D.; Robert, M.; Baffert, K.-A.; Hervé, G.; Quilhot, P.; Attignon, V.; Girod, A.; Chaine, A.; Benassarou, M.; Zrounba, P.; Caux, C.; Ghiringhelli, F.; Lantuejoul, S.; Crozes, C.; Brochériou, I.; Pérol, M.; Fayette, J.; Bertolus, C.; Saintigny, P. Datasets for Gene Expression Profiles of Head and Neck Squamous Cell Carcinoma and Lung Cancer Treated or Not by PD1/PD-L1 Inhibitors. Data Brief 2022, 44, 108556. 10.1016/j.dib.2022.108556.

(49) van den Ende, T.; de Clercq, N. C.; van Berge Henegouwen, M. I.; Gisbertz, S. S.; Geijsen, E. D.; Verhoeven, R. H. A.; Meijer, S. L.; Schokker, S.; Dings, M. P. G.; Bergman, J. J. G. H. M.; Haj Mohammad, N.; Ruurda, J. P.; van Hillegersberg, R.; Mook, S.; Nieuwdorp, M.; de Gruijl, T. D.; Soeratram, T. T. D.; Ylstra, B.; van Grieken, N. C. T.; Bijlsma, M. F.; Hulshof, M. C. C. M.; van Laarhoven, H. W. M. Neoadjuvant Chemoradiotherapy Combined with Atezolizumab for Resectable Esophageal Adenocarcinoma: A Single-Arm Phase II Feasibility Trial (PERFECT). Clin. Cancer Res. 2021, 27 (12), 3351–3359. 10.1158/1078-0432.CCR-20-4443.

(50) Zappasodi, R.; Serganova, I.; Cohen, I. J.; Maeda, M.; Shindo, M.; Senbabaoglu, Y.; Watson, M. J.; Leftin, A.; Maniyar, R.; Verma, S.; Lubin, M.; Ko, M.; Mane, M. M.; Zhong, H.; Liu, C.; Ghosh, A.; Abu-Akeel, M.; Ackerstaff, E.; Koutcher, J. A.; Ho, P.-C.; Delgoffe, G. M.; Blasberg, R.; Wolchok, J. D.; Merghoub, T. CTLA-4 Blockade Drives Loss of Treg Stability in Glycolysis-Low Tumours. Nature 2021, 591 (7851), 652–658. 10.1038/s41586-021-03326-4.

(51) DeVito, N. C.; Sturdivant, M.; Thievanthiran, B.; Xiao, C.; Plebanek, M. P.; Salama, A. K. S.; Beasley, G. M.; Holtzhausen, A.; Novotny-Diermayr, V.; Strickler, J. H.; Hanks, B. A. Pharmacological Wnt Ligand Inhibition Overcomes Key Tumor-Mediated Resistance Pathways to Anti-PD-1 Immunotherapy. Cell Rep. 2021, 35 (5), 109071. 10.1016/j.celrep.2021.109071.

(52) Pusztai, L.; Yau, C.; Wolf, D. M.; Han, H. S.; Du, L.; Wallace, A. M.; String-Reasor, E.; Boughey, J. C.; Chien, A. J.; Elias, A. D.; Beckwith, H.; Nanda, R.; Albain, K. S.; Clark, A. S.; Kemmer, K.; Kalinsky, K.; Isaacs, C.; Thomas, A.; Shatsky, R.; Helsten, T. L.; Forero-Torres, A.; Liu, M. C.; Brown-Swigart, L.; Petricoin, E. F.; Wulfkuhle, J. D.; Asare, S. M.; Wilson, A.; Singhrao, R.; Sit, L.; Hirst, G. L.; Berry, S.; Sanil, A.; Asare, A. L.; Matthews, J. B.; Perlmutter, J.; Melisko, M.; Rugo, H. S.; Schwab, R. B.; Symmans, W. F.; Yee, D.; van’t Veer, L. J.; Hylton, N. M.; DeMichele, A. M.; Berry, D. A.; Esserman, L. J. Durvalumab with Olaparib and Paclitaxel for High-Risk HER2-Negative Stage II/III Breast Cancer: Results from the Adaptively Randomized I-SPY2 Trial. Cancer Cell 2021, 39 (7), 989–998.e5. 10.1016/j.ccell.2021.05.009.

(53) Rose, T. L.; Weir, W. H.; Mayhew, G. M.; Shibata, Y.; Eulitt, P.; Uronis, J. M.; Zhou, M.; Nielsen, M.; Smith, A. B.; Woods, M.; Hayward, M. C.; Salazar, A. H.; Milowsky, M. I.; Wobker, S. E.; McGinty, K.; Millburn, M. V.; Eisner, J. R.; Kim, W. Y. Fibroblast Growth Factor Receptor 3 Alterations and Response to Immune Checkpoint Inhibition in Metastatic Urothelial Cancer: A Real World Experience. Br. J. Cancer 2021, 125 (9), 1251–1260. 10.1038/s41416-021-01488-6.

(54) Mamdani, H.; Schneider, B.; Perkins, S. M.; Burney, H. N.; Kasi, P. M.; Abushahin, L. I.; Birdas, T.; Kesler, K.; Watkins, T. M.; Badve, S. S.; Radovich, M.; Jalal, S. I. A Phase II Trial of Adjuvant Durvalumab Following Trimodality Therapy for Locally Advanced Esophageal and Gastroesophageal Junction Adenocarcinoma: A Big Ten Cancer Research Consortium Study. Front. Oncol. 2021, 11. 10.3389/fonc.2021.736620.

(55) Li, B.; Li, Y.; Zhou, H.; Xu, Y.; Cao, Y.; Cheng, C.; Peng, J.; Li, H.; Zhang, L.; Su, K.; Xu, Z.; Hu, Y.; Lu, J.; Lu, Y.; Qian, L.; Wang, Y.; Zhang, Y.; Liu, Q.; Xie, Y.; Guo, S.; Mehal, W. Z.; Yu, D. Multiomics Identifies Metabolic Subtypes Based on Fatty Acid Degradation Allocating Personalized Treatment in Hepatocellular Carcinoma. Hepatology 2024, 79 (2), 289. 10.1097/HEP.0000000000000553.

(56) Roland, C. L.; Nassif Haddad, E. F.; Keung, E. Z.; Wang, W.-L.; Lazar, A. J.; Lin, H.; Chelvanambi, M.; Parra, E. R.; Wani, K.; Guadagnolo, B. A.; Bishop, A. J.; Burton, E. M.; Hunt, K. K.; Torres, K. E.; Feig, B. W.; Scally, C. P.; Lewis, V. O.; Bird, J. E.; Ratan, R.; Araujo, D.; Zarzour, M. A.; Patel, S.; Benjamin, R.; Conley, A. P.; Livingston, J. A.; Ravi, V.; Tawbi, H. A.; Lin, P. P.; Moon, B. S.; Satcher, R. L.; Mujtaba, B.; Witt, R. G.; Traweek, R. S.; Cope, B.; Lazcano, R.; Wu, C.-C.; Zhou, X.; Mohammad, M. M.; Chu, R. A.; Zhang, J.; Damania, A.; Sahasrabhojane, P.; Tate, T.; Callahan, K.; Nguyen, S.; Ingram, D.; Morey, R.; Crosby, S.; Mathew, G.; Duncan, S.; Lima, C. F.; Blay, J.-Y.; Fridman, W. H.; Shaw, K.; Wistuba, I.; Futreal, A.; Ajami, N.; Wargo, J. A.; Somaiah, N. A Randomized, Non-Comparative Phase 2 Study of Neoadjuvant Immune-Checkpoint Blockade in Retroperitoneal Dedifferentiated Liposarcoma and Extremity/Truncal Undifferentiated Pleomorphic Sarcoma. Nat. Cancer 2024, 5 (4), 625–641. 10.1038/s43018-024-00726-z.

(57) Thibaudin, M.; Fumet, J.-D.; Chibaudel, B.; Bennouna, J.; Borg, C.; Martin-Babau, J.; Cohen, R.; Fonck, M.; Taieb, J.; Limagne, E.; Blanc, J.; Ballot, E.; Hampe, L.; Bon, M.; Daumoine, S.; Peroz, M.; Mananet, H.; Derangère, V.; Boidot, R.; Michaud, H.-A.; Laheurte, C.; Adotevi, O.; Bertaut, A.; Truntzer, C.; Ghiringhelli, F. First-Line Durvalumab and Tremelimumab with Chemotherapy in RAS-Mutated Metastatic Colorectal Cancer: A Phase 1b/2 Trial. Nat. Med. 2023, 29 (8), 2087– 2098. 10.1038/s41591-023-02497-z.

(58) Chin, W. L.; Cook, A. M.; Chee, J.; Principe, N.; Hoang, T. S.; Kidman, J.; Hmon, K. P. W.; Yeow, Y.; Jones, M. E.; Hou, R.; Denisenko, E.; McDonnell, A. M.; Hon, C.-C.; Moody, J.; Anderson, D.; Yip, S.; Cummins, M. M.; Stockler, M. R.; Kok, P.-S.; Brown, C.; John, T.; Kao, S. C.-H.; Karikios, D. J.; O’Byrne, K. J.; Hughes, B. G. M.; Lake, R. A.; Forrest, A. R. R.; Nowak, A. K.; Lassmann, T.; Lesterhuis, W. J. Coupling of Response Biomarkers between Tumor and Peripheral Blood in Patients Undergoing Chemoimmunotherapy. Cell Rep. Med. 2025, 6 (1), 101882. 10.1016/j.xcrm.2024.101882.

(59) Nicolle, R.; Canivet, C.; Palazzo, L.; Napoléon, B.; Ayadi, M.; Pignolet, C.; Cros, J.; Gourgou, S.; Selves, J.; Torrisani, J.; Dusetti, N.; Cordelier, P.; Buscail, L.; Bournet, B.; Bournet, B.; Canivet, C.; Buscail, L.; Carrère, N.; Muscari, F.; Suc, B.; Guimbaud, R.; Couteau, C.; Deslandres, M.; Rivera, P.; Alouany, E.; Fares, N.; Barange, K.; Selves, J.; Gomez-Brouchet, A.; Culetto, A.; Le Cosquer, G.; Jaffrelot, M.; Napoléon, B.; Pujol, B.; Fumex, F.; Desrame, J.; Lefort, C.; Lepilliez, V.; Gincul, R.; Artru, P.; Clavel, L.; Lemaistre, A.-I.; Palazzo, L.; Cros, J.; Tubiana, S.; Flori, N.; Senesse, P.; Colombo, P.-E.; Samalin-Scalzi, E.; Portales, F.; Gourgou, S.; Roca, L.; Ga, C. H.; Plassot, C.; Ychou, M.; Guibert, P.; De La Fouchardière, C.; Sarabi, M.; Peyrat, P.; Tabone-Eglinger, S.; Renard, C.; Piessen, G.; Truant, S.; Saudemont, A.; Millet, G.; Renaud, F.; Leteurtre, E.; Gelé, P.; Assenat, E.; Fabre, J.-M.; Souche, F.-R.; Dupuy, M.; Gorce-Dupuy, A.-M.; Ramos, J.; Seitz, J.-F.; Hardwigsen, J.; Norguet-Monnereau, E.; Grandval, P.; Duluc, M.; Figarella-Branger, D.; Vendrely, V.; Subtil, C.; Terrebonne, E.; Blanc, J.-F.; Merlio, J.-P.; Farges-Bancel, D.; Gornet, J.-M.; Geromin, D.; Vanbiervliet, G.; Frin, A.-C.; Ouvrier, D.; Saint-Paul, M.-C.; Berthélémy, P.; Fouad, C.; Garcia, S.; Lesavre, N.; Gasmi, M.; Barthet, M.; Cottet, V.; Delpierre, C. Predictive Genomic and Transcriptomic Analysis on Endoscopic Ultrasound-Guided Fine Needle Aspiration Materials from Primary Pancreatic Adenocarcinoma: A Prospective Multicentre Study. eBioMedicine 2024, 109, 105373. 10.1016/j.ebiom.2024.105373.

(60) Friedlander, P.; Wassmann, K.; Christenfeld, A. M.; Fisher, D.; Kyi, C.; Kirkwood, J. M.; Bhardwaj, N.; Oh, W. K. Whole-Blood RNA Transcript-Based Models Can Predict Clinical Response in Two Large Independent Clinical Studies of Patients with Advanced Melanoma Treated with the Checkpoint Inhibitor, Tremelimumab. J. Immunother. Cancer 2017, 5 (1). 10.1186/s40425-017-0272-z.

(61) Gide, T. N.; Quek, C.; Menzies, A. M.; Tasker, A. T.; Shang, P.; Holst, J.; Madore, J.; Lim, S. Y.; Velickovic, R.; Wongchenko, M.; Yan, Y.; Lo, S.; Carlino, M. S.; Guminski, A.; Saw, R. P. M.; Pang, A.; McGuire, H. M.; Palendira, U.; Thompson, J. F.; Rizos, H.; Silva, I. P. da; Batten, M.; Scolyer, R. A.; Long, G. V.; Wilmott, J. S. Distinct Immune Cell Populations Define Response to Anti-PD-1 Monotherapy and Anti-PD-1/Anti-CTLA-4 Combined Therapy. Cancer Cell 2019, 35 (2), 238–255.e6. 10.1016/j.ccell.2019.01.003.

(62) Goswami, S.; Gao, J.; Basu, S.; Shapiro, D. D.; Karam, J. A.; Tidwell, R. S.; Ahrar, K.; Campbell, M. T.; Shen, Y.; Trevino, A. E.; Mayer, A. T.; Espejo, A. B.; Seua, C.; Macaluso, M. D.; Chen, Y.; Liu, W.; He, Z.; Yadav, S. S.; Wang, Y.; Rao, P.; Zhao, L.; Zhang, J.; Jindal, S.; Tannir, N. M.; Futreal, A.; Wang, L.; Sharma, P. Immune Checkpoint Inhibitors plus Debulking Surgery for Patients with Metastatic Renal Cell Carcinoma: Clinical Outcomes and Immunological Correlates of a Prospective Pilot Trial. Nat. Commun. 2025, 16 (1), 1846. 10.1038/s41467-025-57009-z.

(63) Hugaboom, M. B.; Wirth, L. V.; Street, K.; Ruthen, N.; Jegede, O. A.; Schindler, N. R.; Shah, V.; Zaemes, J. P.; El Ahmar, N.; Matar, S.; Savla, V.; Choueiri, T. K.; Denize, T.; West, D. J.; McDermott, D. F.; Plimack, E. R.; Sosman, J. A.; Haas, N. B.; Stein, M. N.; Alter, R.; Bilen, M. A.; Hurwitz, M. E.; Hammers, H.; Signoretti, S.; Atkins, M. B.; Wu, C. J.; Braun, D. A. Presence of Tertiary Lymphoid Structures and Exhausted Tissue-Resident T Cells Determines Clinical Response to PD-1 Blockade in Renal Cell Carcinoma. Cancer Discov. 2025, 15 (5), 948–968. 10.1158/2159-8290.CD-24-0991.

(64) Hugo, W.; Zaretsky, J. M.; Sun, L.; Song, C.; Moreno, B. H.; Hu-Lieskovan, S.; Berent-Maoz, B.; Pang, J.; Chmielowski, B.; Cherry, G.; Seja, E.; Lomeli, S.; Kong, X.; Kelley, M. C.; Sosman, J. A.; Johnson, D. B.; Ribas, A.; Lo, R. S. Genomic and Transcriptomic Features of Response to Anti-PD-1 Therapy in Metastatic Melanoma. Cell 2016, 165 (1), 35–44. 10.1016/j.cell.2016.02.065.

(65) Balar, A. V.; Galsky, M. D.; Rosenberg, J. E.; Powles, T.; Petrylak, D. P.; Bellmunt, J.; Loriot, Y.; Necchi, A.; Hoffman-Censits, J.; Perez-Gracia, J. L.; Dawson, N. A.; van der Heijden, M. S.; Dreicer, R.; Srinivas, S.; Retz, M. M.; Joseph, R. W.; Drakaki, A.; Vaishampayan, U. N.; Sridhar, S. S.; Quinn, D. I.; Durán, I.; Shaffer, D. R.; Eigl, B. J.; Grivas, P. D.; Yu, E. Y.; Li, S.; Kadel, E. E.; Boyd, Z.; Bourgon, R.; Hegde, P. S.; Mariathasan, S.; Thåström, A.; Abidoye, O. O.; Fine, G. D.; Bajorin, D. F. Atezolizumab as First-Line Treatment in Cisplatin-Ineligible Patients with Locally Advanced and Metastatic Urothelial Carcinoma: A Single-Arm, Multicentre, Phase 2 Trial. The Lancet 2017, 389 (10064), 67–76. 10.1016/S0140-6736(16)32455-2.

(66) McDermott, D. F.; Huseni, M. A.; Atkins, M. B.; Motzer, R. J.; Rini, B. I.; Escudier, B.; Fong, L.; Joseph, R. W.; Pal, S. K.; Reeves, J. A.; Sznol, M.; Hainsworth, J.; Rathmell, W. K.; Stadler, W. M.; Hutson, T.; Gore, M. E.; Ravaud, A.; Bracarda, S.; Suárez, C.; Danielli, R.; Gruenwald, V.; Choueiri, T. K.; Nickles, D.; Jhunjhunwala, S.; Piault-Louis, E.; Thobhani, A.; Qiu, J.; Chen, D. S.; Hegde, P. S.; Schiff, C.; Fine, G. D.; Powles, T. Clinical Activity and Molecular Correlates of Response to Atezolizumab Alone or in Combination with Bevacizumab versus Sunitinib in Renal Cell Carcinoma. Nat. Med. 2018, 24 (6), 749–757. 10.1038/s41591-018-0053-3.

(67) Motzer, R. J.; Penkov, K.; Haanen, J.; Rini, B.; Albiges, L.; Campbell, M. T.; Venugopal, B.; Kollmannsberger, C.; Negrier, S.; Uemura, M.; Lee, J. L.; Vasiliev, A.; Miller, W. H.; Gurney, H.; Schmidinger, M.; Larkin, J.; Atkins, M. B.; Bedke, J.; Alekseev, B.; Wang, J.; Mariani, M.; Robbins, P. B.; Chudnovsky, A.; Fowst, C.; Hariharan, S.; Huang, B.; Pietro, A. di; Choueiri, T. K. Avelumab plus Axitinib versus Sunitinib for Advanced Renal-Cell Carcinoma. N. Engl. J. Med. 2019, 380 (12), 1103–1115. 10.1056/NEJMoa1816047.

(68) Kim, S. T.; Cristescu, R.; Bass, A. J.; Kim, K.-M.; Odegaard, J. I.; Kim, K.; Liu, X. Q.; Sher, X.; Jung, H.; Lee, M.; Lee, S.; Park, S. H.; Park, J. O.; Park, Y. S.; Lim, H. Y.; Lee, H.; Choi, M.; Talasaz, A.; Kang, P. S.; Cheng, J.; Loboda, A.; Lee, J.; Kang, W. K. Comprehensive Molecular Characterization of Clinical Responses to PD-1 Inhibition in Metastatic Gastric Cancer. Nat. Med. 2018, 24 (9), 1449–1458. 10.1038/s41591-018-0101-z.

(69) Kinget, L.; Naulaerts, S.; Govaerts, J.; Vanmeerbeek, I.; Sprooten, J.; Laureano, R. S.; Dubroja, N.; Shankar, G.; Bosisio, F. M.; Roussel, E.; Verbiest, A.; Finotello, F.; Ausserhofer, M.; Lambrechts, D.; Boeckx, B.; Wozniak, A.; Boon, L.; Kerkhofs, J.; Zucman-Rossi, J.; Albersen, M.; Baldewijns, M.; Beuselinck, B.; Garg, A. D. A Spatial Architecture-Embedding HLA Signature to Predict Clinical Response to Immunotherapy in Renal Cell Carcinoma. Nat. Med. 2024, 30 (6), 1667–1679. 10.1038/s41591-024-02978-9.

(70) Lee, J. S.; Nair, N. U.; Dinstag, G.; Chapman, L.; Chung, Y.; Wang, K.; Sinha, S.; Cha, H.; Kim, D.; Schperberg, A. V.; Srinivasan, A.; Lazar, V.; Rubin, E.; Hwang, S.; Berger, R.; Beker, T.; Ronai, Z.; Hannenhalli, S.; Gilbert, M. R.; Kurzrock, R.; Lee, S.-H.; Aldape, K.; Ruppin, E. Synthetic Lethality-Mediated Precision Oncology via the Tumor Transcriptome. Cell 2021, 184 (9), 2487–2502.e13. 10.1016/j.cell.2021.03.030.

(71) Liu, D.; Schilling, B.; Liu, D.; Sucker, A.; Livingstone, E.; Jerby-Arnon, L.; Zimmer, L.; Gutzmer, R.; Satzger, I.; Loquai, C.; Grabbe, S.; Vokes, N.; Margolis, C. A.; Conway, J.; He, M. X.; Elmarakeby, H.; Dietlein, F.; Miao, D.; Tracy, A.; Gogas, H.; Goldinger, S. M.; Utikal, J.; Blank, C. U.; Rauschenberg, R.; von Bubnoff, D.; Krackhardt, A.; Weide, B.; Haferkamp, S.; Kiecker, F.; Izar, B.; Garraway, L.; Regev, A.; Flaherty, K.; Paschen, A.; Van Allen, E. M.; Schadendorf, D. Integrative Molecular and Clinical Modeling of Clinical Outcomes to PD1 Blockade in Patients with Metastatic Melanoma. Nat. Med. 2019, 25 (12), 1916–1927. 10.1038/s41591-019-0654-5.

(72) Schalper, K. A.; Rodriguez-Ruiz, M. E.; Diez-Valle, R.; López-Janeiro, A.; Porciuncula, A.; Idoate, M. A.; Inogés, S.; de Andrea, C.; López-Diaz de Cerio, A.; Tejada, S.; Berraondo, P.; Villarroel-Espindola, F.; Choi, J.; Gúrpide, A.; Giraldez, M.; Goicoechea, I.; Gallego Perez-Larraya, J.; Sanmamed, M. F.; Perez-Gracia, J. L.; Melero, I. Neoadjuvant Nivolumab Modifies the Tumor Immune Microenvironment in Resectable Glioblastoma. Nat. Med. 2019, 25 (3), 470–476. 10.1038/s41591-018-0339-5.

(73) Nagumo, Y.; Kandori, S.; Kojima, T.; Hamada, K.; Nitta, S.; Chihara, I.; Shiga, M.; Negoro, H.; Mathis, B. J.; Nishiyama, H. Whole-Blood Gene Expression Profiles Correlate with Response to Immune Checkpoint Inhibitors in Patients with Metastatic Renal Cell Carcinoma. Cancers 2022, 14 (24), 6207. 10.3390/cancers14246207.

(74) Nathanson, T.; Ahuja, A.; Rubinsteyn, A.; Aksoy, B. A.; Hellmann, M. D.; Miao, D.; Van Allen, E.; Merghoub, T.; Wolchok, J. D.; Snyder, A.; Hammerbacher, J. Somatic Mutations and Neoepitope Homology in Melanomas Treated with CTLA-4 Blockade. Cancer Immunol. Res. 2017, 5 (1), 84–91. 10.1158/2326-6066.CIR-16-0019.

(75) Galsky, M. D.; Autio, K. A.; Cabanski, C. R.; Wentzel, K.; Graff, J. N.; Friedlander, T. W.; Howes, T. R.; Shotts, K. M.; Densmore, J.; Spasic, M.; Da Silva, D. M.; Chen, R. O.; Lata, J.; Skolnik, J.; Keler, T.; Yellin, M. J.; LaVallee, T. M.; Fairchild, J.; Boffo, S.; O’Donnell-Tormey, J.; Dugan, U.; Bhardwaj, N.; Subudhi, S. K.; Fong, L. Clinical and Translational Results from PORTER, a Multicohort Phase I Platform Trial of Combination Immunotherapy in Metastatic Castration-Resistant Prostate Cancer. Clin. Cancer Res. 2025, 31 (8), 1463–1475. 10.1158/1078-0432.CCR-24-3693.

(76) Padrón, L. J.; Maurer, D. M.; O’Hara, M. H.; O’Reilly, E. M.; Wolff, R. A.; Wainberg, Z. A.; Ko, A. H.; Fisher, G.; Rahma, O.; Lyman, J. P.; Cabanski, C. R.; Yu, J. X.; Pfeiffer, S. M.; Spasic, M.; Xu, J.; Gherardini, P. F.; Karakunnel, J.; Mick, R.; Alanio, C.; Byrne, K. T.; Hollmann, T. J.; Moore, J. S.; Jones, D. D.; Tognetti, M.; Chen, R. O.; Yang, X.; Salvador, L.; Wherry, E. J.; Dugan, U.; O’Donnell-Tormey, J.; Butterfield, L. H.; Hubbard-Lucey, V. M.; Ibrahim, R.; Fairchild, J.; Bucktrout, S.; LaVallee, T. M.; Vonderheide, R. H. Sotigalimab and/or Nivolumab with Chemotherapy in First-Line Metastatic Pancreatic Cancer: Clinical and Immunologic Analyses from the Randomized Phase 2 PRINCE Trial. Nat. Med. 2022, 28 (6), 1167–1177. 10.1038/s41591-022-01829-9.

(77) Cui, C.; Xu, C.; Yang, W.; Chi, Z.; Sheng, X.; Si, L.; Xie, Y.; Yu, J.; Wang, S.; Yu, R.; Guo, J.; Kong, Y. Ratio of the Interferon-γ Signature to the Immunosuppression Signature Predicts Anti-PD-1 Therapy Response in Melanoma. Npj Genomic Med. 2021, 6 (1), 7. 10.1038/s41525-021-00169-w.

(78) Prat, A.; Navarro, A.; Paré, L.; Reguart, N.; Galván, P.; Pascual, T.; Martínez, A.; Nuciforo, P.; Comerma, L.; Alos, L.; Pardo, N.; Cedrés, S.; Fan, C.; Parker, J. S.; Gaba, L.; Victoria, I.; Viñolas, N.; Vivancos, A.; Arance, A.; Felip, E. Immune-Related Gene Expression Profiling After PD-1 Blockade in Non–Small Cell Lung Carcinoma, Head and Neck Squamous Cell Carcinoma, and Melanoma. Cancer Res. 2017, 77 (13), 3540–3550. 10.1158/0008-5472.CAN-16-3556.

(79) Cloughesy, T. F.; Mochizuki, A. Y.; Orpilla, J. R.; Hugo, W.; Lee, A. H.; Davidson, T. B.; Wang, A. C.; Ellingson, B. M.; Rytlewski, J. A.; Sanders, C. M.; Kawaguchi, E. S.; Du, L.; Li, G.; Yong, W. H.; Gaffey, S. C.; Cohen, A. L.; Mellinghoff, I. K.; Lee, E. Q.; Reardon, D. A.; O’Brien, B. J.; Butowski, N. A.; Nghiemphu, P. L.; Clarke, J. L.; Arrillaga-Romany, I. C.; Colman, H.; Kaley, T. J.; de Groot, J. F.; Liau, L. M.; Wen, P. Y.; Prins, R. M. Neoadjuvant Anti-PD-1 Immunotherapy Promotes a Survival Benefit with Intratumoral and Systemic Immune Responses in Recurrent Glioblastoma. Nat. Med. 2019, 25 (3), 477–486. 10.1038/s41591-018-0337-7.

(80) Ravi, A.; Hellmann, M. D.; Arniella, M. B.; Holton, M.; Freeman, S. S.; Naranbhai, V.; Stewart, C.; Leshchiner, I.; Kim, J.; Akiyama, Y.; Griffin, A. T.; Vokes, N. I.; Sakhi, M.; Kamesan, V.; Rizvi, H.; Ricciuti, B.; Forde, P. M.; Anagnostou, V.; Riess, J. W.; Gibbons, D. L.; Pennell, N. A.; Velcheti, V.; Digumarthy, S. R.; Mino-Kenudson, M.; Califano, A.; Heymach, J. V.; Herbst, R. S.; Brahmer, J. R.; Schalper, K. A.; Velculescu, V. E.; Henick, B. S.; Rizvi, N.; Jänne, P. A.; Awad, M. M.; Chow, A.; Greenbaum, B. D.; Luksza, M.; Shaw, A. T.; Wolchok, J.; Hacohen, N.; Getz, G.; Gainor, J. F. Genomic and Transcriptomic Analysis of Checkpoint Blockade Response in Advanced Non-Small Cell Lung Cancer. Nat. Genet. 2023, 55 (5), 807–819. 10.1038/s41588-023-01355-5.

(81) Riaz, N.; Havel, J. J.; Makarov, V.; Desrichard, A.; Urba, W. J.; Sims, J. S.; Hodi, F. S.; Martín-Algarra, S.; Mandal, R.; Sharfman, W. H.; Bhatia, S.; Hwu, W.-J.; Gajewski, T. F.; Slingluff, C. L.; Chowell, D.; Kendall, S. M.; Chang, H.; Shah, R.; Kuo, F.; Morris, L. G. T.; Sidhom, J.-W.; Schneck, J. P.; Horak, C. E.; Weinhold, N.; Chan, T. A. Tumor and Microenvironment Evolution during Immunotherapy with Nivolumab. Cell 2017, 171 (4), 934–949.e16. 10.1016/j.cell.2017.09.028.

(82) Shiuan, E.; Reddy, A.; Dudzinski, S. O.; Lim, A. R.; Sugiura, A.; Hongo, R.; Young, K.; Liu, X.-D.; Smith, C. C.; O’Neal, J.; Dahlman, K. B.; McAlister, R.; Chen, B.; Ruma, K.; Roscoe, N.; Bender, J.; Ward, J.; Kim, J. Y.; Vaupel, C.; Bordeaux, J.; Ganesan, S.; Mayer, T. M.; Riedlinger, G. M.; Vincent, B. G.; Davis, N. B.; Haake, S. M.; Rathmell, J. C.; Jonasch, E.; Rini, B. I.; Rathmell, W. K.; Beckermann, K. E. Clinical Features and Multiplatform Molecular Analysis Assist in Understanding Patient Response to Anti-PD-1/PD-L1 in Renal Cell Carcinoma. Cancers 2021, 13 (6), 1475. 10.3390/cancers13061475.

(83) Subramanian, A.; Nemat-Gorgani, N.; Ellis-Caleo, T. J.; van IJzendoorn, D. G. P.; Sears, T. J.; Somani, A.; Luca, B. A.; Zhou, M. Y.; Bradic, M.; Torres, I. A.; Oladipo, E.; New, C.; Kenney, D. E.; Avedian, R. S.; Steffner, R. J.; Binkley, M. S.; Mohler, D. G.; Tap, W. D.; D’Angelo, S. P.; van de Rijn, M.; Ganjoo, K. N.; Bui, N. Q.; Charville, G. W.; Newman, A. M.; Moding, E. J. Sarcoma Microenvironment Cell States and Ecosystems Are Associated with Prognosis and Predict Response to Immunotherapy. Nat. Cancer 2024, 5 (4), 642–658. 10.1038/s43018-024-00743-y.

(84) Van Allen, E. M.; Miao, D.; Schilling, B.; Shukla, S. A.; Blank, C.; Zimmer, L.; Sucker, A.; Hillen, U.; Geukes Foppen, M. H.; Goldinger, S. M.; Utikal, J.; Hassel, J. C.; Weide, B.; Kaehler, K. C.; Loquai, C.; Mohr, P.; Gutzmer, R.; Dummer, R.; Gabriel, S.; Wu, C. J.; Schadendorf, D.; Garraway, L. A. Genomic Correlates of Response to CTLA-4 Blockade in Metastatic Melanoma. Science 2015, 350 (6257), 207–211. 10.1126/science.aad0095.

(85) Zhao, J.; Chen, A. X.; Gartrell, R. D.; Silverman, A. M.; Aparicio, L.; Chu, T.; Bordbar, D.; Shan, D.; Samanamud, J.; Mahajan, A.; Filip, I.; Orenbuch, R.; Goetz, M.; Yamaguchi, J. T.; Cloney, M.; Horbinski, C.; Lukas, R. V.; Raizer, J.; Rae, A. I.; Yuan, J.; Canoll, P.; Bruce, J. N.; Saenger, Y. M.; Sims, P.; Iwamoto, F. M.; Sonabend, A. M.; Rabadan, R. Immune and Genomic Correlates of Response to Anti-PD-1 Immunotherapy in Glioblastoma. Nat. Med. 2019, 25 (3), 462–469. 10.1038/s41591-019-0349-y.

(86) Wolf, D. M.; Yau, C.; Wulfkuhle, J.; Brown-Swigart, L.; Gallagher, R. I.; Lee, P. R. E.; Zhu, Z.; Magbanua, M. J.; Sayaman, R.; O’Grady, N.; Basu, A.; Delson, A.; Coppé, J. P.; Lu, R.; Braun, J.; Asare, S. M.; Sit, L.; Matthews, J. B.; Perlmutter, J.; Hylton, N.; Liu, M. C.; Pohlmann, P.; Symmans, W. F.; Rugo, H. S.; Isaacs, C.; DeMichele, A. M.; Yee, D.; Berry, D. A.; Pusztai, L.; Petricoin, E. F.; Hirst, G. L.; Esserman, L. J.; van’t Veer, L. J. Redefining Breast Cancer Subtypes to Guide Treatment Prioritization and Maximize Response: Predictive Biomarkers across 10 Cancer Therapies. Cancer Cell 2022, 40 (6), 609–623.e6. 10.1016/j.ccell.2022.05.005.

(87) Liu, D.; Schilling, B.; Liu, D.; Sucker, A.; Livingstone, E.; Jerby-Arnon, L.; Zimmer, L.; Gutzmer, R.; Satzger, I.; Loquai, C.; Grabbe, S.; Vokes, N.; Margolis, C. A.; Conway, J.; He, M. X.; Elmarakeby, H.; Dietlein, F.; Miao, D.; Tracy, A.; Gogas, H.; Goldinger, S. M.; Utikal, J.; Blank, C. U.; Rauschenberg, R.; von Bubnoff, D.; Krackhardt, A.; Weide, B.; Haferkamp, S.; Kiecker, F.; Izar, B.; Garraway, L.; Regev, A.; Flaherty, K.; Paschen, A.; Van Allen, E. M.; Schadendorf, D. Integrative Molecular and Clinical Modeling of Clinical Outcomes to PD1 Blockade in Patients with Metastatic Melanoma. Nat. Med. 2019, 25 (12), 1916–1927. 10.1038/s41591-019-0654-5.

(88) Gérard, A.; Ghiotto, M.; Fos, C.; Guittard, G.; Compagno, D.; Galy, A.; Lemay, S.; Olive, D.; Nunès, J. A. Dok-4 Is a Novel Negative Regulator of T Cell Activation1. J. Immunol. 2009, 182 (12), 7681–7689. 10.4049/jimmunol.0802203.

